# Assessment of Stiffness-Dependent Autophagosome Formation and Apoptosis in Embryonal Rhabdomyosarcoma Tumor Cells

**DOI:** 10.1101/2024.03.01.583012

**Authors:** Serap Sezen, Sevin Adiguzel, Atefeh Zareour, Arezoo Khosravi, Joseph W Gordon, Saeid Ghavami, Ali Zarrabi

## Abstract

Remodeling of the extracellular matrix (ECM) eventually causes the stiffening of tumors and changes to the microenvironment. The stiffening alters the biological processes in cancer cells due to altered signaling through cell surface receptors. Autophagy, a key catabolic process in normal and cancer cells, is thought to be involved in mechano-transduction and the level of autophagy is probably stiffness-dependent. Here, we provide a methodology to study the effect of matrix stiffness on autophagy in embryonal rhabdomyosarcoma cells. To mimic stiffness, we seeded cells on GelMA hydrogel matrices with defined stiffness and evaluated autophagy-related endpoints. We also evaluated autophagy dependent pathways, apoptosis, and cell viability. Specifically, we utilized immunocytochemistry and confocal microscopy to track autophagosome formation through LC3 lipidation. This approach suggests that the use GelMA hydrogels with defined stiffness represent a novel method to evaluate the role of autophagy in embryonal rhabdomyosarcoma and other cancer cells.

**Workflow:** 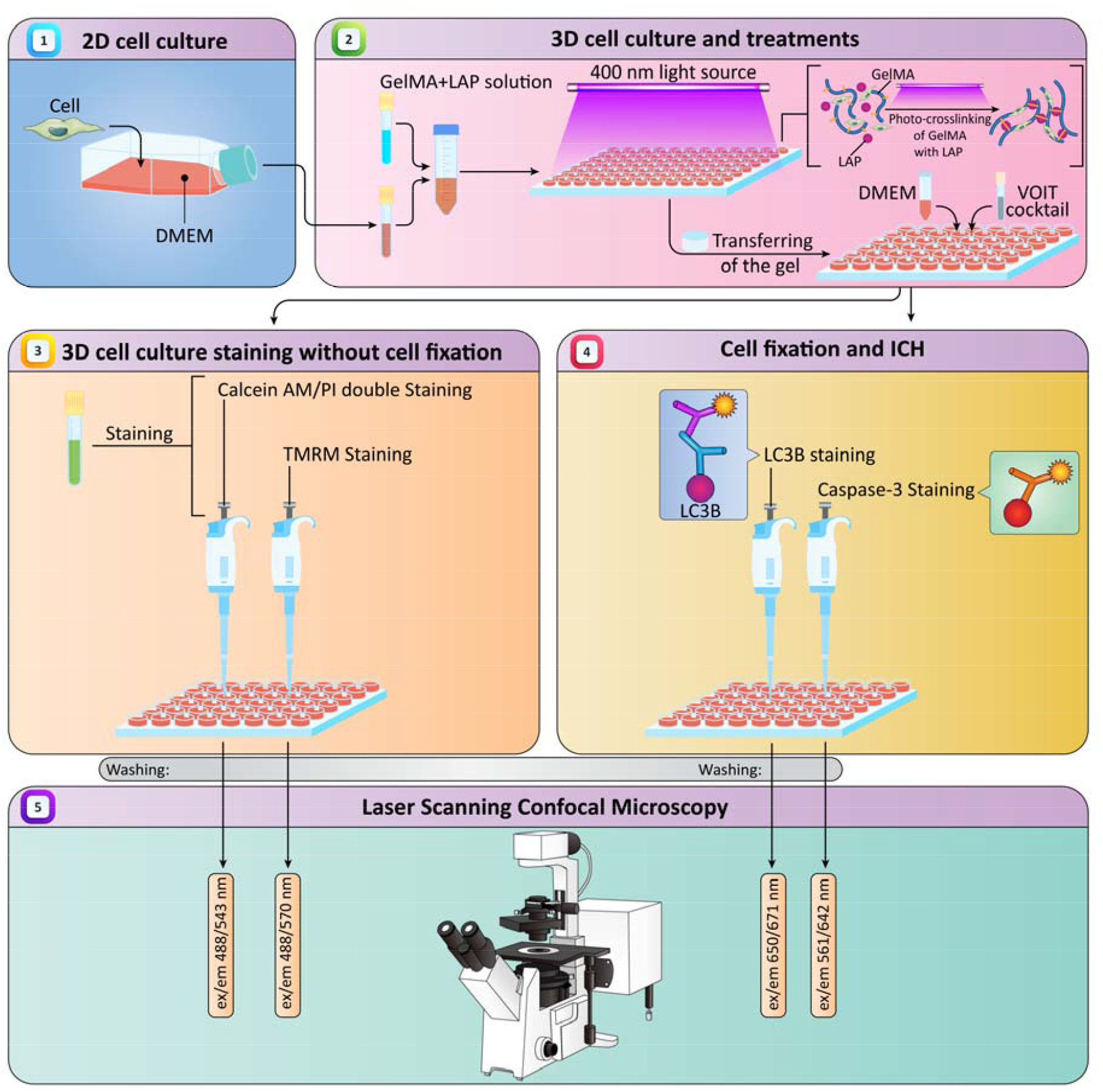

## 1 Introduction

Rhabdomyosarcoma (RMS) is a soft tissue tumor mostly seen in children (70%), and young adults (1). Although RMS is a rare disease, it accounts for 5% of childhood malignancies and 50% of soft tissue sarcomas (2). Two types of RMS are common: alveolar RMS (ARMS) and embryonic RMS (ERMS). ERMS is more prevalent compared to ARMS in both adults and children with wide spectrum of morphology of tumor cells in the clinical findings (2).

During tumor development the biosynthesis of ECM proteins changes resulting in increased tumor stiffness. The increased stiffness of the tumor tissue limits tumor growth but enhances tumor aggressiveness. Through the enzymatic ECM remodeling and degradation, cells can be free from physical barriers favoring tumor growth and invasion, which is a common feature of most cancers (3). The increase in ECM stiffness in solid tumor niche is also associated with the deposition of collagen and lysyl oxidase proteins which eventually results in contraction and deposition, associated with the crosslinking of the stroma. The increase in the stiffness of tumors is considered as a critical step and leads to change in several molecular mechanisms in most cancers. On the contrary, the softening of the tumor ECM slows down the cancer progression and could be considered as targeted therapy for the most tumors (4).

The change in ECM stiffness in tumors also alters autophagy, which is one of the fundamental cellular mechanisms in cancer cells (5,6). Autophagy is a physiological event in eukaryotic cells and plays an important role in cellular hemostasis by eliminating damaged or mis-synthesized proteins as well as dysfunctional organelles, mutant proteins prone to aggregation, and intracellular pathogens (7–9). There are 3 forms of autophagy: chaperone-mediated, microautophagy and macroautophagy (hereafter autophagy) (2,10).

In autophagy, the conversion of microtubule-associated protein 1 light chain 3 (LC3) from the free form (LC3B-I) to the phosphatidyl ethanolamine-conjugated form (LC3B-II) via autophagy-related protein 4 (ATG4) is considered a crucial step for autophagosome formation (11,12). Autophagy is regulated through various stressors that define the fate of the cell (13,14). In particular, it is regulated by cell surface receptors and through mammalian target of rapamycin (mTOR) signaling, which is directly influenced by the ECM stiffness of the tumor niche (15). Moreover, autophagy is also involved in ECM mechano-transduction, so it has been suggested that in normal mammalian cells, autophagy increases with increasing matrix stiffness (16). Besides, matrix stiffness regulates the stromal autophagy and enhances the formation of protumorigenic ECM (17).

The growing tumor cells causes tumor tissue to be up to ten times stiffer than normal tissue (18). ECM stiffness has an impact on the phenotype of malignant cells which is an important parameter often observed in cell-based experiments and drug discovery research to clarify biological pathways. Although *ex vivo* experiments have begun to explore the relationship between ECM stiffness, tumor stiffness and drug activity, a mechanistic-biological link between stiffness and chemotherapy response in RMS has not yet been identified (19). Although tumor stiffness can be monitored in *in vivo*, it is hard to generate tumor with defined stiffness in animal models. The synthesis of hydrogels and their application in tissue engineering has made it possible to develop 3D cell cultures similar to the tumor microenvironment (20,21). The mechanical properties provided by these hydrogels regulate intracellular signaling pathways and studies similar to natural tissues can be completed with such approaches (22). Thus, we developed a method to assess the autophagy based on the matrix stiffness in ERMS cells. We generated cell-laden GelMA hydrogels and measured autophagy using immunocytochemistry techniques in conjunction with confocal microscopy. This approach represents a novel method to evaluate the role of autophagy in embryonal rhabdomyosarcoma and other cancer cells.

## 2 Materials

### 2.1. 2D cell culture

1. Human rhabdoid cell line (A204) (ATCC HTB-82™, Rockville, MD, USA)
2. Mouse muscle cell line (C2C12) (ATCC CRL-1772™ Rockville, MD, USA)
3. Dulbecco’s Modified Eagle’s Medium (DMEM) (PAN BIOTECH™ GmbH, Aidenbach, Germany)
4. Penicillin/Streptomycin (P/S) (PAN BIOTECH™ GmbH, Aidenbach, Germany)
5. Dulbecco’s Phosphate Buffered Saline (DPBS) (PAN BIOTECH™ GmbH, Aidenbach, Germany)
6. Fetal Bovine Serum (FBS) (Biowest, Nuaillé, France)
7. Trypsin-EDTA (PAN BIOTECH™ GmbH, Aidenbach, Germany)

### 2.2. 3D cell culture and treatments

1. Gelatin Methacrylate (GelMA) (CELLINK, Boston, MA, USA)
2. Lithium phenyl-2,4,6-trimethylbenzoylphosphinate (LAP) (Sigma, St. Louis, MO, USA)
3. 96-well and 48-well plates (Greiner BIO ONE, Fisher Scientific, Illkirch, France)
4. Hemocytometer, Counting chambers, Imp. Neubauer (VWR, Radnor, PA, USA)
5. Trypan blue solution 0.4 % (Thermo Fisher Scientific, Wilmington, DE, USA)

### 2.3. 3D cell culture staining

1. Calcein, AM, cell-permeant dye (ThermoFisher Scientific, Wilmington, DE, USA)
2. Propium Iodide Solution (ThermoFisher Scientific, Wilmington, DE, USA)
3. Tetramethylrhodamine methyl ester perchlorate (TMRM) (Sigma Aldrich, St. Louis, MO, USA)
4. Hoechst 33342 (Hellobio, Princeton, NJ, USA)
5. DPBS (PAN BIOTECH™ GmbH, Aidenbach, Germany)
6. DMEM medium (PAN BIOTECH™ GmbH, Aidenbach, Germany)
7. Confocal dish (WVR, Radnor, PA, USA)

### 2.4. Immunocytochemistry (ICC)

1. Paraformaldehyde powder (Sigma–Aldrich Inc. St. Louis, MO, USA)
2. Triton X-100 (Sigma–Aldrich Inc. St. Louis, MO, USA)
3. Bovine Serum Albumin powder (BSA) (Sigma–Aldrich Inc. St. Louis, MO, USA)

8. DPBS (PAN BIOTECH™ GmbH, Aidenbach, Germany)
4. Primary Antibody LC3B, (D11) XP® Rabbit mAb (Cell Signaling Technology, Danvers, MA, USA)
5. Seconder Antibody Alexa Fluor® 488 AffiniPure Donkey Anti-Rabbit IgG (Jackson Immunoresearch)
6. Primary Antibody cleaved Caspase-3 (Asp175) (D3E9) Rabbit mAb (Alexa Fluor® 647 Conjugate) (Cell Signaling Technology, Danvers, MA, USA)
7. Hoechst 33342 (Hellobio, Princeton, NJ, USA)

## 3 Methods

### 3.1. 2D cell culture

1. Prepare complete growth medium including DMEM supplemented with 10% FBS and 1% P/S and store at 4 □C.
2. Obtain A204 and C2C12 cells. Culture both A204 and C2C12 cells with complete medium at 37°C, in a 5% humidified CO_2_ atmosphere.
3. After reaching confluency about 75%, passage cells by removing the medium. Then wash the cells with DPBS. Add 1 ml Trypsin-EDTA solution and incubate for 3 minutes at 37 °C. (**see Note 1**) Add complete medium and pipette to avoid any cell clumps. Collect the cell suspension and transfer to the falcon tube, centrifuge at 300 rpm for 5 minutes. Remove the medium and replace it with fresh medium. Passage the cells 1 to 3.

### 3.2. 3D cell culture

1. Obtain GelMA and LAP. Prepare 5 % w/v and 10 % w/v GelMA solution in 2 ml DPBS at 50 □C under slow stirring in a water bath. Add 0.5 % w/v LAP to GelMA solution and stir for a while. Cover the reaction tube with a piece of aluminum foil.
2. Filter the GelMA+LAP mixture via 0.22 um filter inside the cabinet. Transfer it into a new tube and cover. Keep it at 37 □C water bath until use.
3. Obtain cell pellets by trypsin-EDTA treatment. Count the cell pellets with hemocytometer and trypan blue with standard protocol. Dilute the cell pellets as 1×10^6^ cells per ml of GelMA solution. Transfer the diluted cell pellet into 2 ml Eppendorf tubes.
4. Mix cell pellet with GelMA+LAP solution and pipette slowly. Transfer 50 μl of this solution into the wells of 96-well plate. After dividing this mixture, close the lid of the 96-well plate and apply 400 nm light source for 1 minute for photo-crosslinking of GelMA with LAP (**see Note 2**).
5. After crosslinking complete, transfer cell-laden gels into the wells of the 48-well plate and label them (**see Note 3**). Fill each well of 48-well plate with 250 μl complete DMEM. Grow the cells at 37°C, in a 5% humidified CO_2_ atmosphere overnight before drug treatment.

### 3.3. 3D cell culture staining without cell fixation

#### 3.3.1 Calcein AM/PI double staining

1. Prepare Calcein AM dye and prepare a stock solution of Calcein AM in DMSO at 1 mg/ml. The molarity of this stock solution is 1 mM. This stock solution can be stored for short periods of time at −20 □C.
2. Prepare 1 μM working solution of Calcein AM from 1 mM stock solution in DPBS. This working solution should be prepared freshly.
3. Remove the media and wash the cell-laden gels with DPBS several times.
4. Add Calcein AM working solution just enough to cover the gels. Use aluminum foil to cover the plates.
5. Incubate the cells with Calcein AM working solution for 30 minutes at RT.
6. Remove the Calcein AM solution and wash with DPBS.
7. Obtain 1 mg/ml PI solution. The molarity of this solution is 1.5 mM. Store this so□ution at 4 □C.
8. Prepare 500 nM working solution of PI in DPBS from stock solution.
9. After Calcein AM staining, remove the DPBS and incubate the cell laden gels with PI solution for 5 minutes in dark at RT.
10. Discard the PI solution and wash the gels 3 times with DPBS with 5-minute intervals between each wash. (**See Note 4)**
11. Carefully place the gels on confocal dish and add small amount of DPBS to prevent drying. Observe the cells in laser scanning confocal microscopy. Adjust the parameters for Calcein AM (ex/em 494/517 nm) and PI (ex/em 535/617 nm). Record the images in Z-stack format with 10X objective.

#### 3.3.2 TMRM Staining

1. Obtain TMRM powder. Prepare 5 mg/ml stock solution of TMRM corresponding to 10 mM main stock solution.
2. Dilute 10 mM stock solution into 100 μM stock solution. This stock solution is 1000X of working solution. Both solutions can be stored at −20 □C.
3. Prepare working solution by diluting 1:1000 with DPBS. The working concentration for TMRM is 100 nM. This solution should be prepared freshly.
4. Obtain the treated and control cell-laden gels. Remove the media and wash with DPBS.
5. Add 1X TMRM working solution enough to cover the gels. Cover the plate with aluminum foil and incubate at 37 □C for 30 minutes.
6. Remove the TMRM solution and wash the gels 3 times with DPBS.
7. Carefully place the gels on the confocal dish and add small pieces of DPBS to prevent drying. Adjust the wavelength parameters of TMRM (ex/em 488/570 nm) and objective to 10 X. Start imaging in Z-stack format.

### 3.4. Cell fixation and ICC

#### 3.4.1 LC3B staining

1. Obtain paraformaldehyde, BSA, Triton-X-100 and antibody solutions.
2. Prepare 3.7% formaldehyde solution from powder paraformaldehyde with standard protocol in DPBS. This solution can be stored at −20 □C.
3. Prepare blocking and antibody dilution solutions from DPBS, BSA and Triton-X-100. Blocking solution contains 5% BSA + 0.3% Triton-X in DPBS. Antibody dilution buffer includes 1% BSA and 0.3% Triton-X in DPBS. These solutions can be stored at +4 □C.
4. Fix cells with 3.7% formaldehyde solution at RT for 40 minutes.
5. Discard the formaldehyde solution from wells and wash wells with DPBS 3 times. Allow cells to be washed properly by incubating them for 5 minutes with DPBS in each wash.
6. Add blocking solution enough to cover the gels. Incubate cells with blocking solution for 2 hours at RT.
7. Prepare primary antibody solution containing LC3B, (D11) XP® Rabbit mAb in antibody dilution buffer with a 1:300 dilution. This solution should be prepared freshly (**see Note 5**).
8. Incubate the cells with primary antibody solution at 4 °C overnight. Add antibody solution enough to cover the gels.
9. Remove solutions from wells and wash with DPBS for 3 times with 5-min intervals between each wash.
10. Prepare secondary antibody solution containing Alexa Fluor® 488 AffiniPure Donkey Anti-Rabbit IgG in antibody dilution buffer at a 1:300 dilution ratio.
11. Add secondary antibody solution just enough to cover the gels and incubate with seconder antibody solution for 4 h in dark at RT.
12. Discard secondary antibody solution and wash 3 times with DPBS with a 5-minute incubation in each wash.
13. Obtain Hoechst 33342 (**see Note 5**). Prepare 1 mg/ml Hoechst solution in DMSO. This stock can be stored at −20 □C for months. Prepare working solution of Hoechst by diluting stock into 1 μg/ml with DPBS.
14. Add Hoechst working solution to wells enough to cover the gels and allow to incubate for 30 minutes at RT in dark.
15. Remove Hoechst solution and wash 3 times with DPBS with a 5-minute incubation in each wash.
16. Adjust the filters of confocal microscopy for visualization. For LC3B, Alexa Fluor® 488 AffiniPure Donkey Anti-Rabbit IgG (ex/em 495/519 nm) will be imagined with defined wavelengths.

#### 3.4.2. Caspase-3 Staining

1. Perform cell fixation as indicated above.
2. Use the same antibody dilution buffer.
3. Obtain cleaved Caspase-3 (Asp175) (D3E9) Rabbit mAb (Alexa Fluor® 647 Conjugate). This antibody is conjugated with florescence marker and does not require seconder antibody for labelling.
4. Prepare primary antibody solutions containing cleaved Caspase-3 (Asp175) (D3E9) Rabbit mAb (Alexa Fluor® 647 Conjugate) in antibody dilution buffer with a 1:300 dilution. This solution should be prepared freshly.
5. Incubate the cells with primary antibody solutions at 4 °C overnight. Add antibody solutions enough to cover the gels. Cover the plate with aluminum foil.
6. Remove solutions from wells and wash with DPBS for 3 times with 5-min intervals between each wash.
7. Adjust the filters of confocal microscopy for visualization of cleaved Caspase-3 (Asp175) (D3E9) Rabbit mAb (Alexa Fluor® 647 Conjugate) (ex/em 650/671 nm).

### 3.5. Laser Scanning Confocal Microscopy

1. Turn on the device and program (**see Note 6**).
2. In the program, adjust the parameters including objective (**see Note 7**) and lasers (**Table 1**).
3. First place the gel containing confocal dish and observe the cells with one laser channel by coarse and fine adjustment. Adjust acquisition parameters including laser intensity, detector gain and pinhole. Add another channel when needed for multiple imaging.
4. Create a Z-Stack method. Start imaging the slices.
5. Process the data.

### 3.4 Interpretation of the results

The viability of the cells was first studied with Calcein AM/PI double staining. This allows us to visualize the effect of matrix stiffness on the viability of the cells. **Figure 1**. shows the result of Calcein AM/PI double staining of A204 cells grown for 48 hours on 5-GelMA and 10-GelMA matrices. **Figure 1**. also represents the results of TMRM staining. TMRM indicates the mitochondrial membrane potential of the cells, which has direct relationship with the apoptosis levels of the cells.

**Table 1.** Laser excitation and emission wavelengths for florescence indicators.

| Indicator | Excitation | Emission |
| --- | --- | --- |
| Calcein AM | 488 | 543 |
| PI | 561 | 642 |
| TMRM | 561 | 642 |
| Alexa Fluor™<br>488 | 488 | 543 |
| Alexa Fluor™<br>647 | 561 | 642 |
| Hoechst 33342 | 405 | 642 |

**Figure 1.**
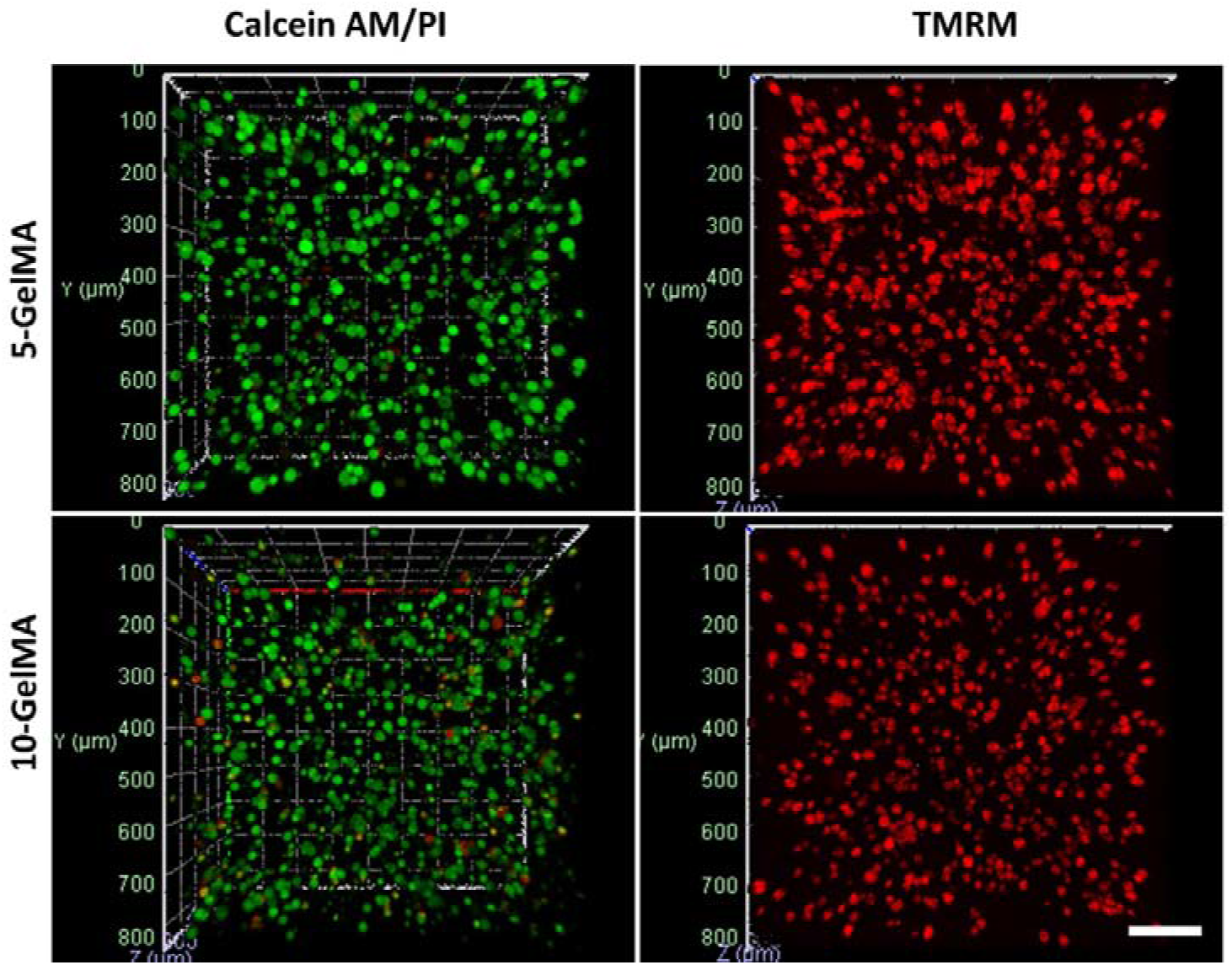
Confocal images of A204 cells seeded on 5-GelMA and 10-GelMA matrices for 48 hours stained with Calcein AM/PI double staining (left) and TMRM (right). The images are taken via 10X objective and represented as Z-Stack format. Scale bar: 100 μm.

For apoptosis, the level of cleaved Caspase-3 was also imaged with ICC technique. **Figure 2**. shows the confocal images of A204 cells seeded on 5-GelMA and 10-GelMA matrices and grown for 48 hours.

**Figure 2.**
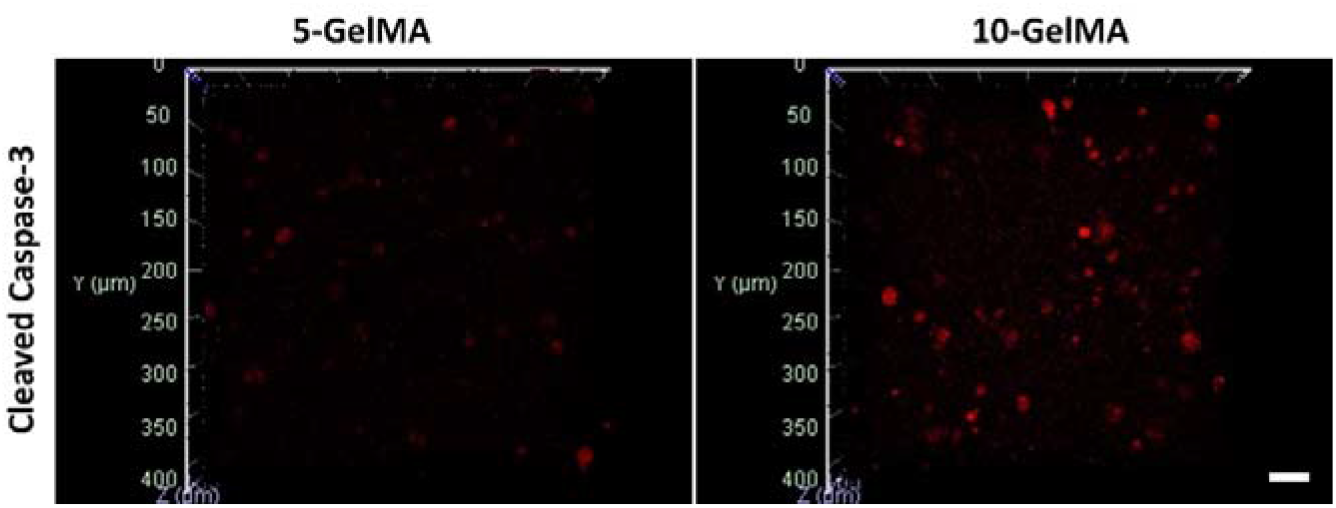
Confocal images of A204 cells seeded on 5-GelMA and 10-GelMA matrices for 48 hours. Cells were fixed and stained for cleaved Caspase-3 with ICC. The images are taken via 20X objective and represented as Z-Stack format. Scale bar: 50 μm.

In the following section, autophagosome was studied. The LC3B puncta, and their altered levels were imagined with ICC, followed by nuclear staining. **Figure 3**. shows ICC of A204 cells seeded on 5-GelMA and 10-GelMA matrices.

**Figure 3.**
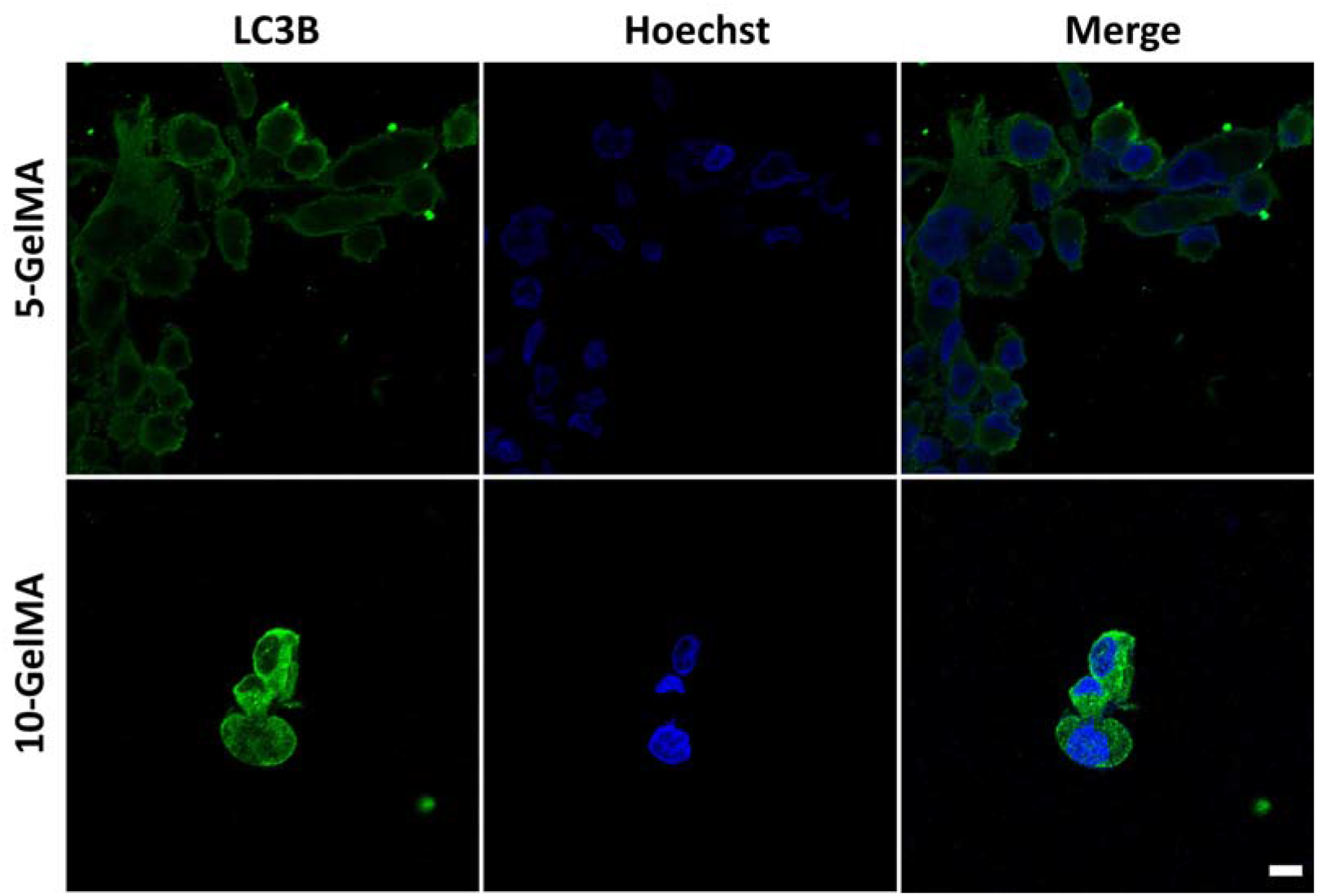
Confocal images of A204 cells after ICC staining. Cells were seeded on 5-GelMA and 10-GelMA matrices and grown for 48 hours. The images were taken with 63X objective with oil. Scale bar: 10 μm.

## Conflict of Interest

The authors declare that the research was conducted in the absence of any commercial or financial relationships that could be construed as a potential conflict of interest.

## Acknowledgement

SG and AZ conceptualized and designed the study. SZ performed the experiments. SG and SZ analyzed the data. SZ and S Kh wrote the manuscript. SZ, SG, AZ, SA, JWG, and AKh reviewed and edited the manuscript. AZ funded and SG supervised the experiments.

## Funding

This study was funded in part by URGP (University of Manitoba) (SG) and 1002-short term R&D funding program (221Z107) (AZ).

## Notes

### NOTE 1

The amount of the Trypsin-EDTA and incubation time is given for the cells seeded to the T25 flask. The amount and time can change according to the flask size.

### NOTE 2

GelMA+LAP mixture should be kept in dark in order to prevent pre-gelation.

### NOTE 3

Once the gelation is complete, the transparent solution turns into blurry gel. These gels are brittle and should be carefully transferred from the 96-wells to the 48-wells filled with the medium. If the gels are dried before transferring, small drop of DMEM medium can be added to keep the gels wet.

### NOTE 4

Before each imaging, the gels should be kept in DPBS. For stained samples, the plates are covered with aluminum foil to prevent photobleaching under light. The imaging should be completed at the same day when staining and ICC protocols are completed.

### NOTE 5

In this study, we used Hoechst 33342 dye. This dye is cell permeant nuclear counterstain. Instead of this one, DAPI can also be used. DAPI can be suitable for fixed cells, although Hoechst can stain both live and dead cells.

### NOTE 6

As scanning laser confocal microscopy, we used LSM 710, Axio Observer (Carl Zeiss, Oberkochen, Germany) equipped with Zeiss Zen software (version 2010). For each microscopy, the given parameters should be altered to get better images.

### NOTE 7

For 10X imaging, EC Plan-Neofluar 10x/0.30 M27 objective was set. For 63X imaging, we used oil-based objective Plan-Apochromat 63x / 1.40 Oil DIC M27.

## Notes

### Competing Interest Statement

The authors have declared no competing interest.

## References

1. Ramadan F, Fahs A, Ghayad SE, Saab R. Signaling pathways in Rhabdomyosarcoma invasion and metastasis. Cancer Metastasis Rev [Internet]. 2020;39(1):287–301. Available from: 10.1007/s10555-020-09860-3

2. Zarrabi A, Perrin D, Kavoosi M, Sommer M, Sezen S, Mehrbod P, et al. Rhabdomyosarcoma: current therapy, challenges, and future approaches to treatment strategies. Cancers (Basel). 2023;15(21):5269.

3. Gonzalez-Molina J, Kirchhof KM, Rathod B, Moyano-Galceran L, Calvo-Noriega M, Kokaraki G, et al. Mechanical Confinement and DDR1 Signaling Synergize to Regulate Collagen-Induced Apoptosis in Rhabdomyosarcoma Cells. Adv Sci [Internet]. 2022 Oct 1;9(28):2202552. Available from: 10.1002/advs.202202552

4. Liang R, Song G. Matrix stiffness-driven cancer progression and the targeted therapeutic strategy. Mechanobiol Med [Internet]. 2023;1(2):100013. Available from: https://www.sciencedirect.com/science/article/pii/S294990702300013X

5. Liang P, Wang B. An autophagy-independent role of ULK1/ULK2 in mechanotransduction and breast cancer cell migration. Autophagy [Internet].:1–2. Available from: 10.1080/15548627.2023.2300916

6. Gong T, Wu D, Pan H, Sun Z, Yao X, Wang D, et al. Biomimetic Microenvironmental Stiffness Boosts Stemness of Pancreatic Ductal Adenocarcinoma via Augmented Autophagy. ACS Biomater Sci Eng [Internet]. 2023 Sep 11;9(9):5347–60. Available from: 10.1021/acsbiomaterials.3c00487

7. Behrooz AB, Cordani M, Donadelli M, Ghavami S. Metastatic outgrowth via the two-way interplay of autophagy and metabolism. Biochim Biophys Acta (BBA)-Molecular Basis Dis. 2023;166824.

8. Siapoush S, Rezaei R, Alavifard H, Hatami B, Zali MR, Vosough M, et al. Therapeutic implications of targeting autophagy and TGF-β crosstalk for the treatment of liver fibrosis. Life Sci. 2023;121894.

9. Alizadeh J, Kochan MM, Stewart VD, Drewnik DA, Hannila SS, Ghavami S. Inhibition of autophagy flux promotes secretion of chondroitin sulfate proteoglycans in primary rat astrocytes. Mol Neurobiol. 2021;58(12):6077–91.

10. Parzych KR, Klionsky DJ. An overview of autophagy: morphology, mechanism, and regulation. Antioxid Redox Signal [Internet]. 2013/08/02. 2014 Jan 20;20(3):460–73. Available from: https://pubmed.ncbi.nlm.nih.gov/23725295

11. Alizadeh J, Kavoosi M, Singh N, Lorzadeh S, Ravandi A, Kidane B, et al. Regulation of Autophagy via Carbohydrate and Lipid Metabolism in Cancer. Cancers (Basel). 2023;15(8):2195.

12. Hajiahmadi S, Lorzadeh S, Iranpour R, Karima S, Rajabibazl M, Shahsavari Z, et al. Temozolomide, simvastatin and acetylshikonin combination induces mitochondrial-dependent apoptosis in GBM cells, which is regulated by autophagy. Biology (Basel). 2023;12(2):302.

13. Martelli A, Omrani M, Zarghooni M, Citi V, Brogi S, Calderone V, et al. New Visions on Natural Products and Cancer Therapy: Autophagy and Related Regulatory Pathways. Cancers (Basel). 2022;14(23):5839.

14. Dalvand A, da Silva Rosa SC, Ghavami S, Marzban H. Potential role of TGFB and autophagy in early crebellum development. Biochem Biophys Reports. 2022;32:101358.

15. D’Agostino S, Tombolan L, Saggioro M, Frasson C, Rampazzo E, Pellegrini S, et al. Rhabdomyosarcoma Cells Produce Their Own Extracellular Matrix With Minimal Involvement of Cancer-Associated Fibroblasts: A Preliminary Study. Front Oncol [Internet]. 2021 Jan 29;10:600980. Available from: https://pubmed.ncbi.nlm.nih.gov/33585217

16. Li Y, Randriantsilefisoa R, Chen J, Cuellar-Camacho JL, Liang W, Li W. Matrix Stiffness Regulates Chemosensitivity, Stemness Characteristics, and Autophagy in Breast Cancer Cells. ACS Appl Bio Mater [Internet]. 2020 Jul 20;3(7):4474–85. Available from: 10.1021/acsabm.0c00448

17. Hupfer A, Brichkina A, Koeniger A, Keber C, Denkert C, Pfefferle P, et al. Matrix stiffness drives stromal autophagy and promotes formation of a protumorigenic niche. Proc Natl Acad Sci [Internet]. 2021 Oct 5;118(40):e2105367118. Available from: 10.1073/pnas.2105367118

18. Lu P, Weaver VM, Werb Z. The extracellular matrix: a dynamic niche in cancer progression. J Cell Biol [Internet]. 2012 Feb 20;196(4):395–406. Available from: https://pubmed.ncbi.nlm.nih.gov/22351925

19. Schrader J, Gordon-Walker TT, Aucott RL, van Deemter M, Quaas A, Walsh S, et al. Matrix stiffness modulates proliferation, chemotherapeutic response, and dormancy in hepatocellular carcinoma cells. Hepatology [Internet]. 2011 Apr;53(4):1192–205. Available from: https://pubmed.ncbi.nlm.nih.gov/21442631

20. Annabi N, Tamayol A, Uquillas JA, Akbari M, Bertassoni LE, Cha C, et al. 25th Anniversary Article: Rational Design and Applications of Hydrogels in Regenerative Medicine. Adv Mater [Internet]. 2014 Jan 1;26(1):85–124. Available from: 10.1002/adma.201303233

21. Stefanek E, Samiei E, Kavoosi M, Esmaeillou M, Roustai Geraylow K, Emami A, et al. A bioengineering method for modeling alveolar Rhabdomyosarcoma and assessing chemotherapy responses. MethodsX [Internet]. 2021;8:101473. Available from: https://www.sciencedirect.com/science/article/pii/S2215016121002661

22. tibbitt MW, Anseth KS. Hydrogels as extracellular matrix mimics for 3D cell culture. Biotechnol Bioeng [Internet]. 2009 Jul 1;103(4):655–63. Available from: https://pubmed.ncbi.nlm.nih.gov/19472329

